# DNA methylation analysis of amplicons from individuals exposed to maternal tobacco use during pregnancy, and offspring conduct problems in childhood and adolescence

**DOI:** 10.1101/2020.07.02.183285

**Authors:** Alexandra J. Noble, John F. Pearson, Alasdair D. Noble, Joseph M. Boden, L. John Horwood, Martin A. Kennedy, Amy J. Osborne

## Abstract

Maternal tobacco smoking during pregnancy is a large driver of health inequalities and a higher prevalence of conduct problem has been observed in exposed offspring. Further, maternal tobacco use during pregnancy can also alter offspring DNA methylation. However, currently, limited molecular evidence have been found to support this observation. Thus we aim to examine the association between maternal tobacco use in pregnancy and whether offspring Conduct problems is mediated by tobacco exposure-induced via DNA methylation differences. Understanding the etiology of the causal link will be crucial in the early identification and treatment of CP in children and adolescents. DNA was sourced from the Christchurch Health and Development Study, a longitudinal birth cohort studied for over 40 years in New Zealand. Bisulfite-based amplicon sequencing of 10 loci known to play a role in neurodevelopment, or with associations with CP phenotypes, was undertaken. We identified nominally significant differential DNA methylation at specific CpG sites in *CYP1A1, ASH2L* and *MEF2C* in individuals with Conduct problems who were exposed to tobacco *in utero.* We conclude that environmentally-induced DNA methylation differences could play a role in the observed link between maternal tobacco use during pregnancy and childhood/adolescent Conduct problems However, larger sample sizes are needed to produce an adequate amount of power to investigate this interaction further.

## Introduction

Mothers who smoked tobacco during pregnancy have a higher prevalence of offspring developing a conduct disorder phenotype compared to mothers who did not smoke (Wakschlag et al. 1997). This association has been proven in several different cohort studies and the observation have remained following adjustment for various other confounding factors, for example, socio economic status, maternal age, substance abuse, parental anti-social personality, and maladaptive parenting (Wakschlag et al. 1997; Joelsson et al. 2016). However, there is limited molecular evidence to suggest a link between *in utero* tobacco exposure and offspring conduct disorder, thus a direct link between *in utero* tobacco exposure and Conduct problems (CP) remains elusive. Previously we conducted a pilot study assessing differential DNA methylation in a small cohort of individuals who were exposed to tobacco *in utero,* with sub-groups of individuals defined as having high conduct disorder scores (Noble et al. 2021). We found nominally significant DNA methylation changes with this interaction in several genes associated with neurodevelopment (cite epic in utero paper). Due to the limitations of using a small sample size combined with an array containing a large number of loci, results were underpowered, therefore observations were unable to reach genome wide significance. However, the biological relevance of these nominally significant CpG loci to the CP phenotype, combined with further research which has suggested an epigenetic link between *in utero* tobacco exposure and ADHD (Sengupta et al. 2017), implies that the link between DNA methylation and CP development in tobacco-exposed offspring should be investigated more fully.

Here, we will further pursue this hypothesis, by exploring differential methylation in genes that have known roles during *in utero* neurodevelopment and CP phenotypes, to understand whether DNA methylation may help explain the relationship between *in utero* tobacco exposure and development of CP in offspring. We applied a targeted approach via bisulfite-based amplicon sequencing (BSAS) of regions of genes involved in neurodevelopment. Amplicon sequencing has the ability to interrogate a region of the genome, therefore specifically targeting consecutive CpG sites in a row. We then assessed differential methylation in the DNA of participants from the Christchurch Health and Development Study (CHDS) whose mothers consumed tobacco during pregnancy, with high and low CP scores, and compared this to controls who were not exposed. This approach allows us to specifically ask whether DNA methylation at genes involved in neurodevelopment and CP phenotypes are specifically differentially methylated in the DNA of offspring with CP, who were exposed to tobacco *in utero.* A significant interaction here would provide further support for a role for DNA methylation in the link between *in utero* exposure and CP development, something which has so far proved elusive.

## Methods

### Sample

A sub-group of individuals from the CHDS were selected for this study (Table 1). This longitudinal study originally included 97% of all the children (N= 1265) born in the Christchurch, New Zealand urban region during a period in mid-1977 and has been studied at 24 time points from birth to age 40 (N= 987 at age 30). All participants were aged between 28-30 when blood samples and DNA was extracted.

**Table 1.** CHDS subsets selected for analysis. The range of conduct problem scores in each category is indicated in brackets. A score of 53 or more is the top quartile for CP, a score of 60 or more the top decile for CP.

|  | Group 1 | Group 2 | Group 3 |
| --- | --- | --- | --- |
|  | Exposed *in utero* and never smokers | Exposed *in utero* and a regular smoker | Not exposed *in utero* and never smokers |
|  | n= 32 | n= 32 | n= 32 |
| Sex |  |  |  |
| Male | 69% | 72% | 60% |
| Female | 31% | 28% | 40% |
| Tobacco smoking status at the time of blood collection |  |  |  |
| Never | 100% | 0% | 100% |
| Regular | 0% | 100% | 0% |
| Conduct problem score (CPS) |  |  |  |
| Low CPS (<46) | n= 16 (42-46) | n= 16 (42-46) | n= 16 (41-43) |
| High CPS (>53) | n= 16 (53-75) | n= 16 (60-85) | n= 16 (53-68) |

For the subsets studied in this report, CHDS participants were chosen based on their *in utero* tobacco exposure status, their adult smoking status, and their CP scores. Group 1 consisted of individuals who were exposed *in utero* to tobacco smoke, and never smokers at the time blood samples were taken (N= 32). Group 2 consisted of individuals who were exposed *in utero* to tobacco smoke and were themselves regular smokers at the time the blood was taken (N= 32). Group 3 consisted of individuals who were not exposed to tobacco *in utero,* and never smokers at the time blood was taken (N= 32). *In utero* tobacco exposure was defined as 10+ cigarettes per day throughout pregnancy. Within each group of 32, 16 individuals were selected with a ‘high’ score on a measure of childhood CP at age 7-9 years and 16 with a ‘low’ score. Severity of childhood CP was assessed using an instrument that combined selected items from the Rutter and Conners child behaviour checklists (Rutter M 1970; Conners 1970, 1969; Fergusson, Horwood, and Lloyd 1991) as completed by parents and teachers at annual intervals from 7-9 years. Parental and teacher reports were summed and averaged over the three years (Fergusson, Horwood, and Ridder 2005) to derive a robust scale measure of the extent to which the child exhibited conduct disordered/oppositional behaviours (mean (SD)= 50.1(7.9); range 41-97). For the purposes of this report a ‘high’ score was defined as falling into the top quartile of the score distribution (scores >53) and a ‘low’ score was defined as scores < 46.

### Bisulfite-based amplicon sequencing

Bisulfite-based amplicon sequencing (BSAS) was carried out as described (Noble et al. 2020). DNA was extracted from whole blood samples using the Kingfisher Flex System (Thermo Scientific, Waltham, MA USA). DNA was quantified via nanodrop (Thermo Scientific, Waltham, MA USA). Bisulfite treatment was carried out using the EZ DNA Methylation-Gold kit (Zymo Research, Irvine, CA, USA) as per the manufacturer’s instructions. DNA samples were then diluted to a final concentration of 100 ng/μl.

Amplicons for sequencing (Table 2 and Supplementary Table 1) were picked based upon several criteria: i) previously published differential DNA methylation in response to *in utero* tobacco smoking; ii) known associations with *in utero* brain development, and; iii) known associations with CP phenotypes. Primers were then designed to flank the CpG sites of interest, ~350 base pairs (bp) in total, or to amplify the promoter region of the gene if a specific CpG site was not known. Multiple pairs of primers were designed to amplify larger regions.

**Table 2.**
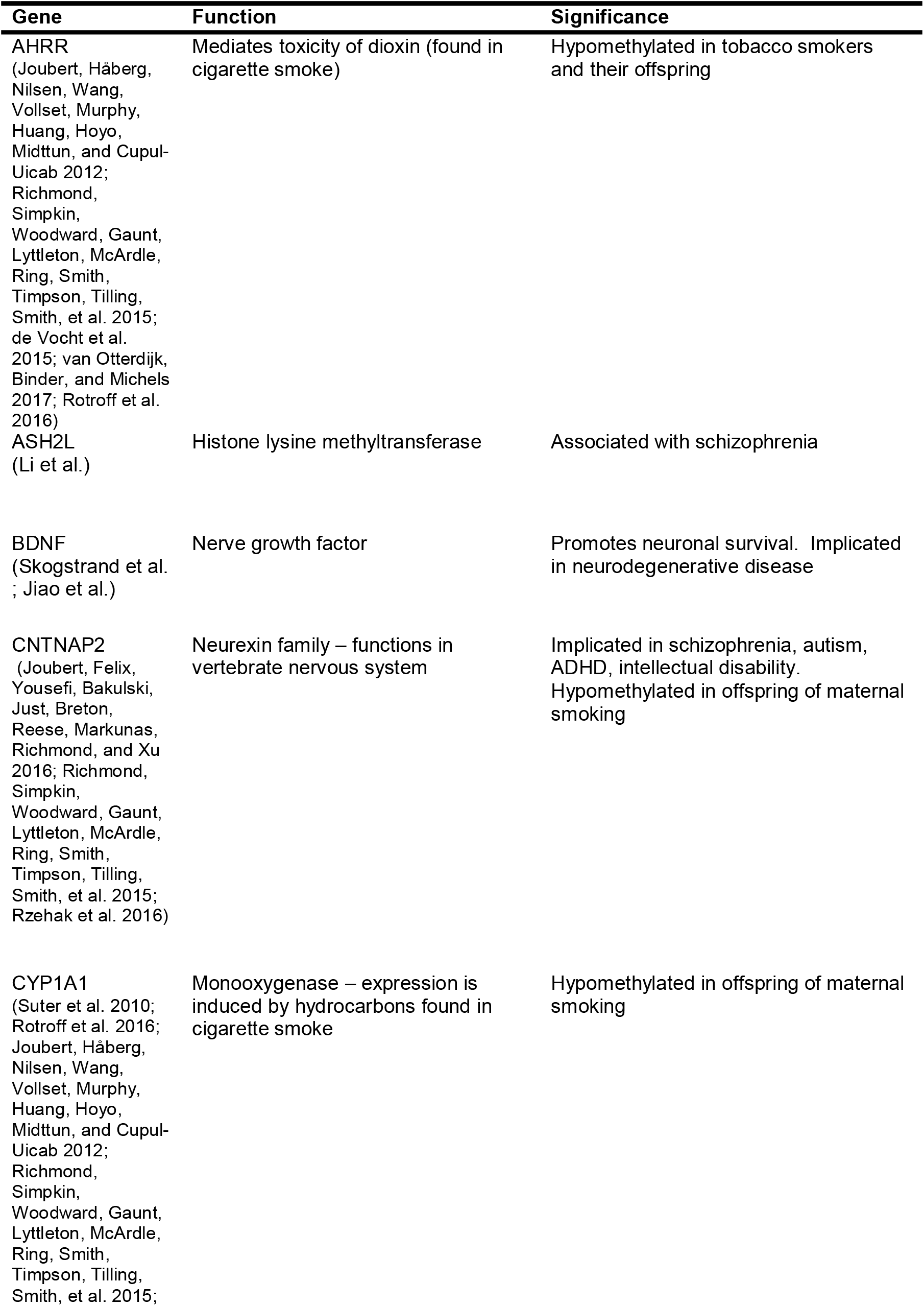
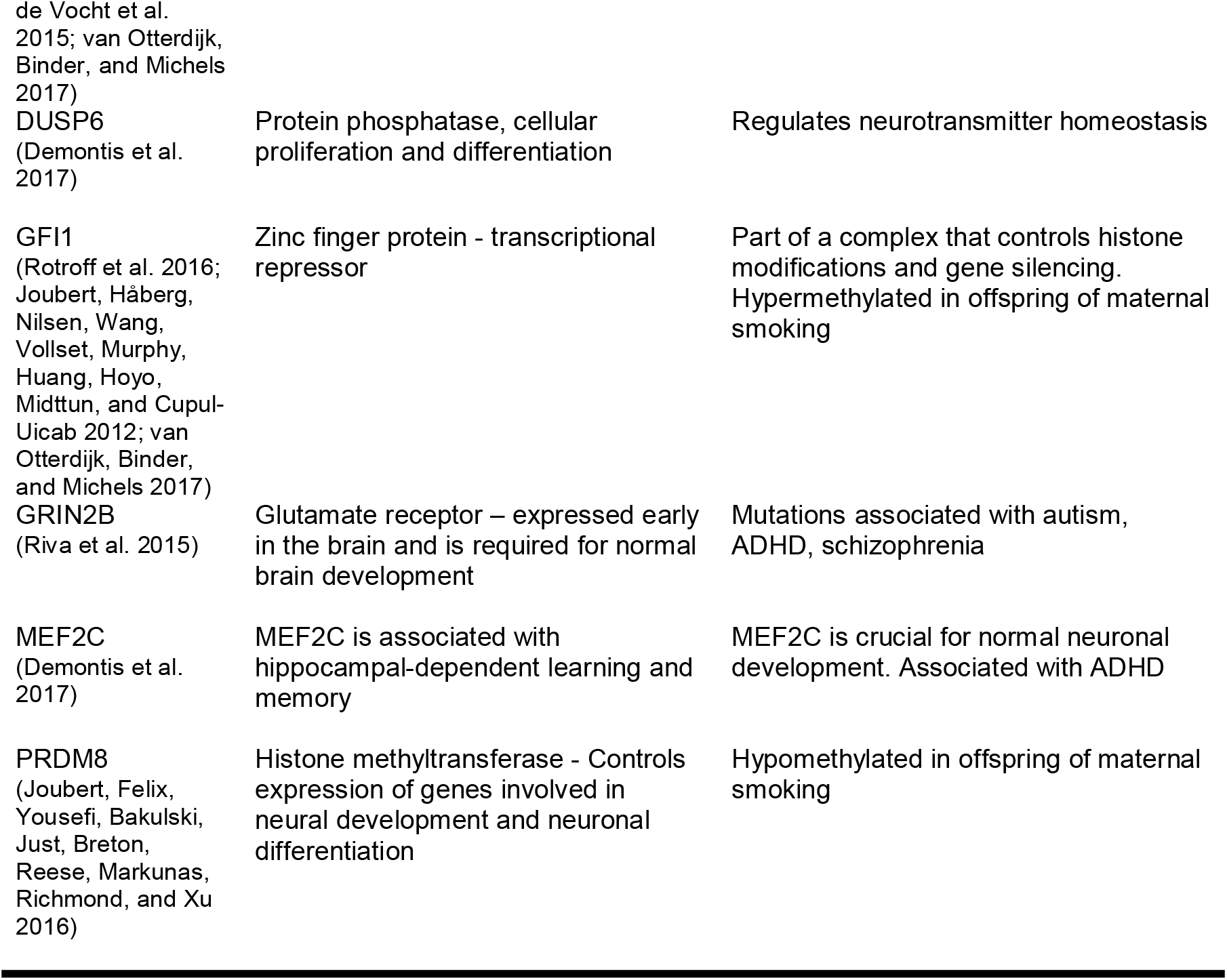
Genes selected to investigate the link between in utero tobacco exposure and CP.

Bisulfite-converted DNA was amplified via PCR, using KAPA Taq HotStart DNA Polymerase (Sigma, Aldrich) under the following conditions: 95 °C for 10 min, 95 °C for 30 sec, 59 °C for 20 sec, 72 °C for 7 min, and held at 4 C° using the Mastercycler Nexus (Eppendorf, Australia). This was then cycled a total of 40 times. PCR products were purified with the Zymo DNA Clean & Concentrator Kit™ (Zymo Research, Irvine, CA, USA).

Following PCR, DNA was cleaned up with Agencourt^®^ AMPure^®^ XP beads (Beckman Coulter) and washed with 80% ethanol and allowed to air-dry. DNA was then eluted with 52.5 μl of 10 mM Tris pH 8.5 before being placed back into the magnetic stand. Once the supernatant had cleared, 50 μl was aliquoted for the experiment. DNA samples were quantified using the Quant-iT™ PicoGreen™ dsDNA Assay kit (Thermo Fisher) using the FLUROstar^®^ Omega (BMG Labtech).

Samples were processed using the Illumina MiSeq™ 500 cycle Kit V2 and sequenced on the Illumina MiSeq™ system by Massey Genome Service (Palmerston North). Illumina MiSeq™ sequences were trimmed using SolexaQA++ software (Cox, Peterson, and Biggs 2010) and aligned to FASTA bisulfite converted reference sequences using the package Bowtie2 (version 2.3.4.3) Each individual read was then aligned to all reference sequences using the methylation-specific package Bismark (Krueger and Andrews 2011).

### Statistics

Differential DNA methylation was assessed using the package edgeR (Chen et al.). Coverage level was set to greater or equal to ‘‘8’’ across unmethylated and methylated counts, as recommended by (Chen et al.). Two models were used - the first was a bivariate model, to assess differences between the *in utero* exposed to tobacco compared to the non-exposed control group (model 1).

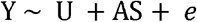

The second was a multiple regression to assess the interaction term *in utero* maternal smoke exposure and offspring conduct problem score (high or low, model 2).

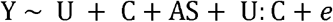

Where, *Y* is defined as the methylation M ratio, *U* is the exposed/unexposed *in utero* to maternal smoking, *C* is conduct problem score with high conduct problem score < 53 and low conduct problem core < 46, *AS* adult smoker/non-smoker and *e* is the unexplained variation or error tem.

This model was fitted with both ANOVA parameters and with contrasts between *in utero* exposure groups (exposed - non-exposed) within CP score levels. Top tables were constructed using the topTags function in edgeR, Log fold change, average log counts per million, and in some cases F statistic and were calculated and nominal significance was given for P < 0.05, these were then corrected using FDR. Scatter plots with the inclusion of confidence intervals were constructed from log transformed normalised methylated and unmethylated counts.

## Results

Here we assessed DNA methylation within 10 separate genes (Table 2). DNA sequence data for 15 amplicons from these 10 genes (Supplementary Table 1) was generated, comprising a total of 280 CpG sites. These CpG sites included a combination of sites previously identified as differentially methylated, as well as amplification of all CpGs within the promoter region of genes associated with *in utero* neurodevelopment and CP phenotypes (Table 2). Differential methylation across these CpG sites was calculated to address whether any were specifically differentially methylated in individuals with CP, in response to *in utero* tobacco exposure.

### Quantification of DNA methylation at previously reported CpG sites in response to *in utero* exposure to tobacco

Initially, we attempted to validate in our cohort (age ~28-30 years) five CpG sites which have been previously reported to be differentially methylated in the DNA of cord blood from newborns, and whole blood from children and adolescents (ages newborn to 17) in response to *in utero* tobacco exposure (Table 1). Data were partitioned into those individuals exposed *in utero,* and those who were not (model 1), to assess whether or not BSAS could detect previously reported CpG sites (Table 3).

**Table 3.**
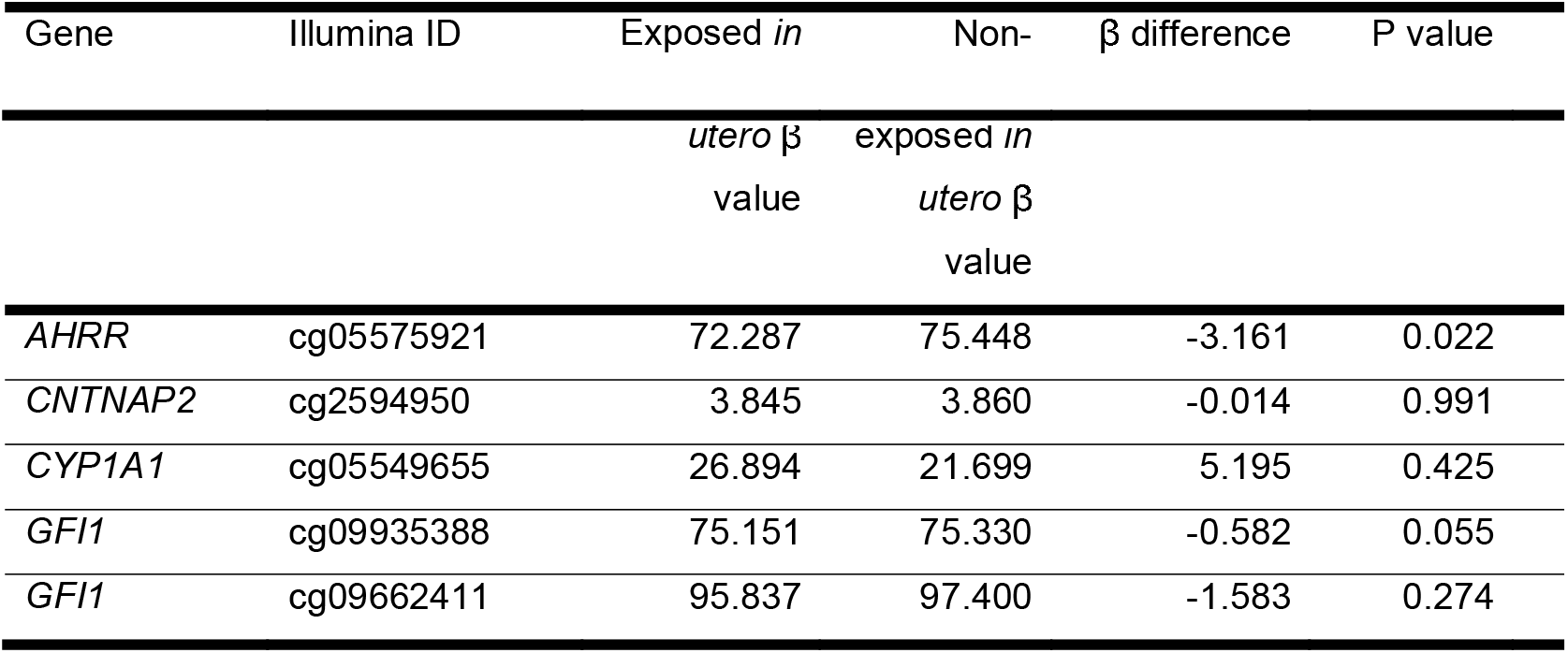
Previously reported CpG sites showing differential DNA methylation in response to *in utero* tobacco exposure, and their average methylation values in individuals from this cohort.

| Gene | Illumina ID | Exposed *in* | Non- | $\beta$ difference | P value |
| --- | --- | --- | --- | --- | --- |

| | | *utero* $\beta$<br>value | exposed *in*<br>*utero* $\beta$<br>value | | |
| --- | --- | --- | --- | --- | --- |
| *AHRR* | cg05575921 | 72.287 | 75.448 | -3.161 | 0.022 |
| *CNTNAP2* | cg2594950 | 3.845 | 3.860 | -0.014 | 0.991 |
| *CYP1A1* | cg05549655 | 26.894 | 21.699 | 5.195 | 0.425 |
| *GFI1* | cg09935388 | 75.151 | 75.330 | -0.582 | 0.055 |
| *GFI1* | cg09662411 | 95.837 | 97.400 | -1.583 | 0.274 |

*AHRR* (cg05575921) displayed a 3.1% decrease in DNA methylation between exposed and non-exposed individuals, at a nominal P value of 0.02. This site has been previously identified as hypomethylated in adult tobacco smokers, as well as in postnatal cord blood samples between *in utero* tobacco-exposed and non-exposed individuals. The probe cg05549655 in the gene *CYP1A1* displayed a 5.19% increase in DNA methylation in the *in utero* exposed group, however, this site did not reach nominal statistical significance in our cohort. Cg09935388 and cg09662411 in *GFI1* were unable to be replicated as differentially methylated between the exposed and the non-exposed groups (no significant change in β values). Both CpG sites show hypomethylation, supporting previous observations of differential methylation within this gene. *CNTNAP2* (cg2594950) was similarly unable to be validated in our cohort using the method BSAS.

### Differentially methylated CpGs under the interaction of *in utero* tobacco exposure and CP

Differential methylation dependent on both *in utero* exposure and CP score was found at 10 loci in six genes at nominal significance level, none were significant after correcting for false discovery rate (Table 4).

**Table 4.**
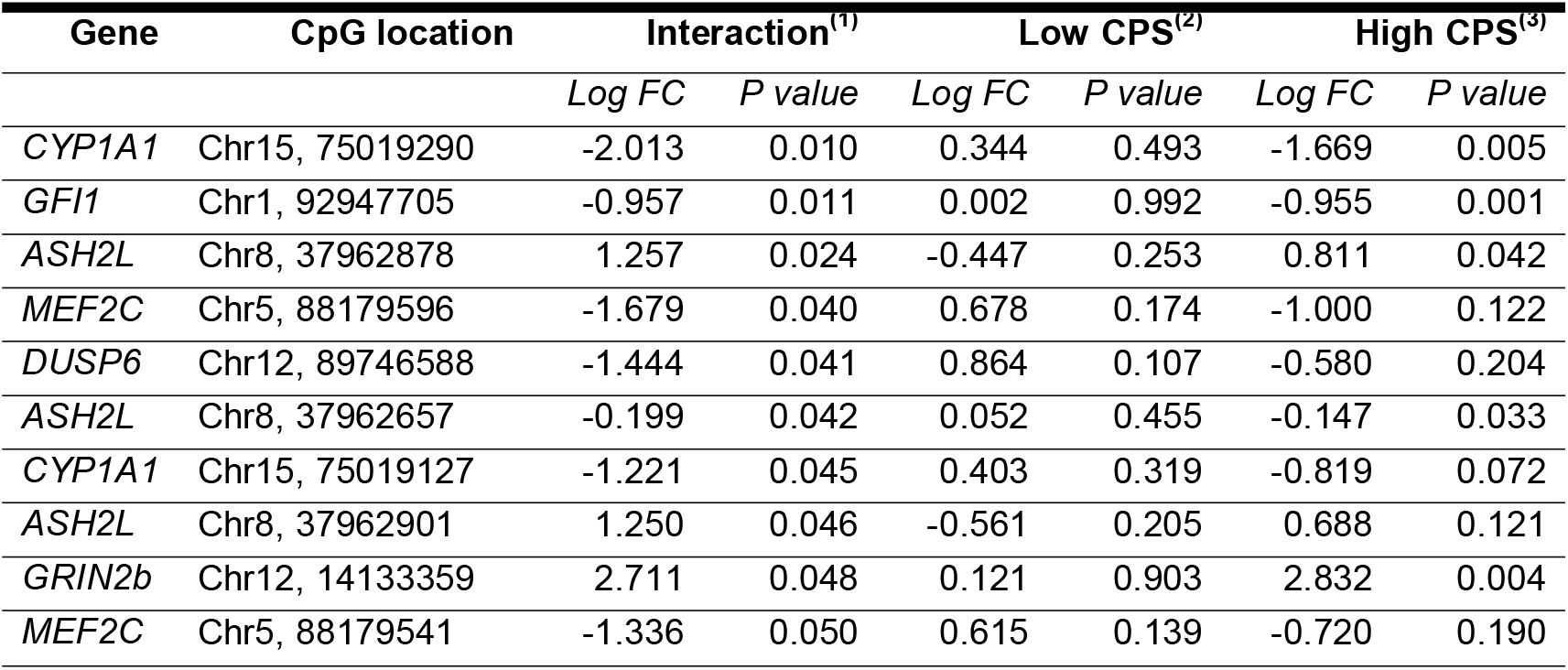
CpG sites where differential methylation between conduct problem scores differs with *in utero* exposure at P< 0.05. Log Fold Change (FC) and P values (unadjusted) from log ratio tests for the effect on normalized methylation ratios of: (1) interaction between *in utero* exposure and Conduct Problem Score, (2) *In utero* exposed - non-exposed contrast within Low CPS and (3) within High CPS participants. Loci with nominally significant (P< 0.05) interaction shown, all FDR P values > 0.05.

Of these CpG sites, five of the 10 CpG sites were found in the following genes: *CYP1A1, GFI1,* ASH2L *and GRIN2b.* Differential methylation between *in utero* exposed and non-exposed associated with for high conduct scores. No nominal significance from the interaction observed in association with low conduct scores. The top three CpG sites with nominal significance under the interaction are displayed in Figure 1. Here, differential methylation is found in response to high CP score and no differences are seen between the exposed and non-exposed low CP groups (Figure 1).

## Discussion

*In utero* tobacco exposure is known to alter DNA methylation at the genome-wide level in offspring (Joubert, Haberg, et al. 2012; Joubert, Felix, Yousefi, Bakulski, Just, Breton, Reese, Markunas, Richmond, Xu, et al. 2016) (Joubert Bonnie et al. 2012; Richmond et al. 2014). The later-life implications of these tobacco-induced DNA methylation changes are unclear, however, an association between *in utero* tobacco exposure and CP has previously been observed (Sengupta et al. 2017). Given the complex etiology of CP phenotypes (Acosta, Arcos-Burgos, and Muenke 2004; Beaver et al. 2007; Salvatore and Dick 2018) and the vast array of socioeconomic variables associated with tobacco use (Lantz et al. 1998), proving a causal link between maternal smoking and offspring CP is inherently challenging. Previously we quantified tobacco-induced DNA methylation changes that associate with CP phenotypes in offspring exposed to tobacco *in utero* (via maternal smoking) using the Illumina EPIC array, with results indicating that methylation was altered at genes that may have roles in neurodevelopment and CP phenotypes. However, due to a combination of a comparatively small sample size relative to the number of loci on the array, only nominal significance was observed. The data suggested a role for DNA methylation in the link between exposure and CP, so here we chose a panel of genes with known roles in these things, to see if methylation is changed at these loci too. Due to our small sample size we need to try an alternative approach. We’ve shown that BSAS previously is very good for targeting differential methylated so here we use BSAS at some targeted genes to see if we can detect differential methylation specific to the interaction between high CP score and exposure?

### Validation of previously identified differentially methylated CpG from *in utero* tobacco exposure

First, we asked whether differentially methylated CpGs that have been previously associated with *in utero* tobacco exposure were supported by this cohort. Here, we present validation of differential methylation of a CpG site within the gene *AHRR* (cg05575921). *AHRR* is a well-defined tobacco smoking gene, which is consistently represented in tobacco methylation data. *AHRR* has previously been found to be differentially methylated in response to *in utero* tobacco exposure (Richmond, Simpkin, Woodward, Gaunt, Lyttleton, McArdle, Ring, Smith, Timpson, Tilling, Davey Smith, et al. 2015; Joubert, Håberg, Nilsen, Wang, Vollset, Murphy, Huang, Hoyo, Midttun, Cupul-Uicab, et al. 2012; de Vocht et al. 2015). Four other CpG sites investigated here due to previous association with *in utero* tobacco exposure were not differentially methylated in our data. However, the direction of methylation change was supported at all five sites investigated (Tehranifar et al. 2018; Rauschert et al. 2019; Rotroff et al. 2016). We suggest that further investigation in a larger cohort may lead to nominal significance at the sites in *CYP1A1, CNTNAP2,* and *GFI1.*

### Identification of *in utero* exposure-related differentially methylated CpG sites that are specific to individuals with high CP scores

Epidemiological data suggests that there is an increased association between *in utero* tobacco exposure and behavioural disorder in children and adolescents (Carter et al. 2008; Mick et al. 2002). Thus, here, we investigated DNA methylation changes induced by *in utero* tobacco exposure as a potential molecular mechanism of dysfunction that could link the phenotypic trait of CP to maternal tobacco use during pregnancy. We therefore analysed DNA methylation patterns within our gene panel in response to *in utero* tobacco exposure and its interaction with CP status. A total of 10 CpG sites in seven genes were found to display nominal significance in DNA methylation in response to *in utero* tobacco exposure and CP in this cohort (Table 4).

In the 10 CpG sites we identified under the interaction, *CYP1A1* showed greater magnitude differential methylation in high CP scores (exposed *in utero* vs. non-exposed with high CPS), with reduced reversed or no evidence of differential methylation at the same sites with low CP score. This indicates that within the observed nominal methylation changes the interaction was being driven in the high CP score group. One gene *(ASH2L),* contained three nominally significantly differentially methylated CpG sites, and *CYP1A1* and *MEF2C* both had two.

*CYP1A1* (Cytochrome P450 family 1 subfamily A member 1) is a well-established marker for *in utero* tobacco smoke exposure (Richmond et al. 2018; Lee Ken et al. 2015; Richmond et al. 2014; Tehranifar et al. 2018). Neither of the two sites we observed under the interaction have probes at these locations on the Illumina array system, thus emphasising a benefit of amplicon sequencing compared to an array-based method. Variant differences in *CYP1A1* have previously been associated with child behavioural problems at age 2, from prenatal maternal environmental tobacco smoke (Hsieh et al. 2010). This highlights the need for this gene to be further investigated for its role in the development of conduct problems following *in utero* tobacco exposure.

Three CpG sites from the gene *ASH2L* (ASH2 like histone lysine methyltransferase complex subunit) showed in consistent levels of differential methylation in response to *in utero* tobacco exposure and CP, with two displaying hyper- and one hypomethylation. *ASH2L* has been found to interact with *MEF2C* (Myocyte enhancer factor 2C) to mediate changes in histone 3 lysine 4 trimethylation (H3K4me3) (Jung et al.). Here, we detected nominal significance at two CpG sites within *MEF2C* (chr5, 88179596 and 88179541). Both of these sites were associated with a greater level of hypomethylation in participants who were exposed to tobacco *in utero* with high CP scores in this cohort, although not at the FDR significance level. *MEF2C* plays a role in neural crest formation during development, where tissue-specific inactivation of the gene results in embryonic lethality (Verzi et al. 2007). Further, *MEF2* interacts with oxytocin, which is affiliated with prosocial behaviours (Kosfeld et al. 2005; Zak, Stanton, and Ahmadi 2007). Alterations to oxytocin have been shown to change the morphology of neurons via *MEF2A* (Meyer et al. 2018; Meyer et al. 2020). Functional roles of the gene in relation to early neuronal development still remain unclear, however it is thought to play a crucial role (Harrington et al. 2016). Recent research in animal models suggests that nicotine-dependent induction of the *ASH2L* and *MEF2C* complex during development induces alterations that could lead to fundamental changes in the brain (Jung et al.).

While we cannot assert causality, our targeted approach shows that *in utero* tobacco exposure may be altering methylation at CpG sites associated with neural phenotypes which persist into adulthood and are then associated with increased risk of high CP.

## Conclusion

Here we have presented preliminary data to suggest that the association between maternal tobacco use during pregnancy and the development of CP in children and adolescents may in part be mediated by altered DNA methylation, induced by *in utero* tobacco exposure during development, at genes that have roles in *in utero* brain development and CP phenotypes. We acknowledge the limitations of this study described above, however, the data presented here are suggestive of a role for DNA methylation in the link between *in utero* tobacco exposure and offspring CP. Our findings should stimulate further study using larger sample sizes.

## Abbreviations

CP: Conduct problems
CHDS: Christchurch health and development study
BSAS: Bisulfite based amplicon sequencing
SIDS: Sudden infant death syndrome
ADHD: Attention-deficit hyperactivity disorder
DOHaD: Developmental origins of human health and disease
GFI1: Growth Factor Independent one transcriptional repressor
CPS: Conduct disorder score
AHRR: Aryl hydrocarbon receptor repressor
ASH2L: ASH2 like histone lysine methyltransferase complex subunit
BDNF: Brain-derived neurotrophic factor,
CNTNAP2: Contactin associated protein 2
CYP1A1: Cytochrome P450 Family 1 Subfamily A Member 1
DUSP6: Dual specificity phosphatase 6
GRIN2b: Glutamate Ionotropic Receptor NMDA Type Subunit 2B
MEF2C: Myocyte enhancer factor 2C
PRDM8: PR/SET Domain 8
FC: Fold change
CPM: Counts per million
FDR: False discovery rate

## Funding

Funding for this study came from the Maurice and Phyllis Paykel Trust. CHDS was funded by the Health Research Council of New Zealand (Programme Grant 16/600). CMRF supplied funding for the manuscript to be written.

## Availability of data

Upon request.

## Contributions

AJN-molecular lab work, data analysis, and major contributor to manuscript. JFP-study design, data analysis, and major contributor to manuscript. ADN-data analysis JMB and LJH study design, provided DNA samples via CHDS. MAK-study design and over view. AJO-study design, molecular lab work, major contributor to manuscript and source of funding. All authors read and approved the final manuscript.

## Acknowledgements

Not applicable

## Ethics declarations

All aspects of the study were approved by the Southern Health and Disability Ethics Committee, under application number CTB/04/11/234/AM10 “Collection of DNA in the Christchurch Health and Development Study”.

## Consent for publication

Not applicable

## Competing interests

The authors declare that they have no competing interests.

## Notes

### Competing Interest Statement

The authors have declared no competing interest.

